# General markers of conscious visual perception and their timing

**DOI:** 10.1101/008995

**Authors:** Renate Rutiku, Jaan Aru, Annika Tallinn, Talis Bachmann

**Affiliations:** Institute of Psychology, University of Tartu, Näituse 2, Tartu 50409, Estonia; Institute of Public Law, University of Tartu (Tallinn branch), Teatri väljak 3-207, Tallinn 10143, Estonia; Institute of Computer Science, University of Tartu, J. Liivi 2, Tartu 50409, Estonia

**Keywords:** NCC, ERP, denoising, VAN, P300

## Abstract

The goal of the present investigation was to identify reliable markers of conscious visual perception and to characterize their onset latency and its variability. To that end many visual stimuli from different categories were presented at near-threshold contrast and contrastive analyses were carried out on 100 balanced subsets of the data. N200 and P300 were the two reliable markers of conscious perception common to all perceived stimuli and absent for all non-perceived stimuli. The estimated mean onset latency for both markers was shortly after 200 ms. However, the onset latency of both of these markers of conscious perception showed considerable variability depending on which subsets of the data were considered. Some of this variability could be attributed to noise, but it was first and foremost the amplitude fluctuation in the condition without conscious perception that explained the variability in onset latencies of the markers of conscious perception. The present results help to understand why different studies have observed different onset times for the neural correlates of conscious perception. Moreover, the consciousness markers explored here have more generality as stimulus specificity was reduced.

## 1. Introduction

How long does it take from the moment when a stimulus is presented in the environment until the conscious experience of the stimulus starts to arise? Despite the decades-long quest for the neural correlates of consciousness (NCC) it is not known **at what time after stimulus onset they occur**. Some results suggest that conscious perception is a relatively late process (e.g. Sergent et al., 2005; Del Cul et al., 2007). Others point to the importance of mid-latency markers (Koivisto & Revonsuo, 2010). Still others have found very early correlates for conscious perception (e.g. Pins & Ffytche, 2003; Aru & Bachmann, 2009).

The reasons for these discrepancies are still largely unknown, but it is plausible to assume that procedural differences between studies are one major factor. A wide variety of different paradigms, stimulus material, recording conditions etc. have been used to identify NCC (see Koivisto & Revonsuo (2010) for an overview). This makes studies inherently difficult to compare. For example, if experiments employ restricted categories of stimuli it is hard to tell whether the resulting NCC are markers of only one category or whether they can be generalized to other categories as well. It is also known that the latency of processes correlating with consciousness may shift as much as 100 ms depending on stimulus predictability (Melloni et al., 2011). If the stimulus set of a study consists of only a few items then perceptual events inevitably become more predictable and the accompanying consequences on NCC latencies would have to be quantified. Consequently, it is not surprising that even studies that have found very similar NCC tend to report varying onset latencies of these markers.

Despite this unavoidable degree of incomparability between studies, it would still be important to know how reliable NCC are within one study and how variable their timing is. Surprisingly, it has not yet been thoroughly characterized how much NCC vary when only the data from one experiment are considered. To bring out the relevance of this question, one could first consider how NCC are typically identified. The identification of NCC is most often based on contrastive analysis (Aru et al., 2012). One searches for markers that are uniquely present or reliably more strongly present in the averaged activity of the condition where a stimulus was consciously perceived compared to the condition where a stimulus was not consciously perceived. It is important to note that the reliability and onset latency of the resulting markers can be influenced by a number of different reasons.

For example, it is possible that the latency of the true NCC shifts from trial to trial. This would spread out the averaged activity in the condition with conscious perception and the mean onset latency of NCC would become less accurate. Navajas et al. (2013) have demonstrated a similar effect for the face-sensitive N170 component if stimulus uncertainty is increased due to added noise. Results from a contrastive analysis may also be influenced by factors not directly related to the NCC proper. Different noise profiles may accompany the signal in different trials. Again, this would influence the onset latency of NCC. One assumes that task irrelevant noise is mostly averaged out when means are created over trials, but this is of course not completely true. Random noise summation will contribute somewhat also to averaged ERPs leading to at least a small effect and thus also on the onset of statistical differences between conditions.

To make matters worse, one cannot even be sure that it is only the signal and noise profiles of the condition with conscious perception that dictate NCC reliability and onset latency. The above described concerns apply to the condition without conscious perception as well. This is because for delineating NCC, trials with conscious perception are compared against those without conscious perception of the target. Only the significant differences are considered as candidates of NCC (Aru et al., 2012), but the timing of these significant differences also depends on the trials in the condition without conscious perception. For characterizing the latency of the NCC it would be necessary to know how much each of these factors contributes to the results of a contrastive analysis in order to arrive at the best estimate of the true NCC.

The present study is designed to address the above described issues. We employed an experimental paradigm where the role of visual categorical restriction and stimulus predictability are reduced. To that end we use many different stimuli with varying characteristics and we present these stimuli on perceptual threshold. We hypothesize that **for the described paradigm there is at least one marker of conscious perception common to all perceived stimuli and absent for all non-perceived stimuli.** We call this the general marker of NCC, **gmNCC** in short. Our first goal is to find which EEG correlates qualify as gmNCC in our experimental paradigm. Our second goal is to study any possible variability in the onset latency of gmNCC and to characterize the causes of this variability as thoroughly as possible. We recognize that the strength of EEG lies in its temporal precision and thus, as a method, it is first and foremost suited for finding answers to questions about timing. Thus, we want to offer a **precise estimate of gmNCC onset latency** that can be used in further research. We hope that our research sheds light on the question why different studies have come up with different onset times for the correlates of conscious perception.

## 2. Materials and Methods

### 2.1. Subjects

22 subjects participated in the EEG experiment. All subjects were healthy and had normal or corrected-to-normal vision. Data from 4 subjects were not included in the analyses due to a high number of noisy electrodes or too many trials with artifacts. The remaining 18 subjects (8 male) were 18 – 31 years old (mean = 23.2, median = 22, SD = 3.6). 1 subject was left-handed. All the included subjects had more than 49 trials in each condition (m = 101, median = 102, SD = 22.5) and less than 6.3% of interpolated data (m = 2.6%, SD = 1.8%). All subjects gave written informed consent prior to participation and received monetary compensation as a reward. The study was approved by the ethics committee of University of Tartu and the experiment was undertaken in compliance with national legislation and the Declaration of Helsinki.

### 2.2. Stimuli

The stimulus set consisted of 70 monochrome drawings. The drawings depicted objects from 6 different categories. 4 categories were further divided into line-drawings and solid forms. Thus, there were 10 different types of stimuli: line-drawings of graphical figures, solid graphical figures, short words, line-drawings of man-made objects, solid forms of man-made objects, line-drawings of faces, line-drawings of animated nature, solid forms of animated nature, line-drawings of inanimate nature, solid forms of inanimate nature. Figure 1 depicts examples for each of the different stimulus types. All stimuli were collected from online databases. Occasionally, stimuli were edited manually to keep the number of filled pixels i.e. the contrast energy comparable for all solid forms including text and all line-drawings including faces.

**Figure 1.**
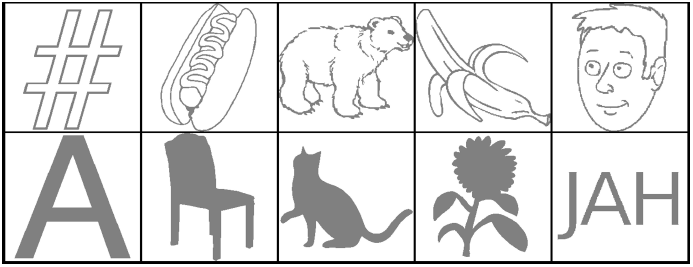
Examples for all ten stimulus types. The contrast of the stimuli was changed to render them near-threshold, thus turning them more lighter or darker.

In order to display stimuli at perceptual threshold their contrast has to be accordingly low. Not all of our stimuli have the same threshold contrast, however. An earlier pilot experiment indicated that for the present stimulus set there are 5 groups of stimuli with roughly similar threshold contrasts within each group: text, solid graphical figures, line-drawings of graphical figures, solid forms of all other figures and line-drawings of all other figures. Thus, contrast was adjusted separately for each of these 5 groups.

The 5 appropriate contrasts were determined with the help of a short pre-experiment prior to the main experiment. The pre-experiment was very similar to the main experiment (see 2.3 Task and design), except that a separate set of stimuli including all the 10 stimulus types was used. Each stimulus (19 in total) was presented twice on 4 adjacent contrast levels. The specific contrast levels were different for each of the 5 contrast groups. They were typical threshold contrasts for these groups as indicated by an earlier pilot experiment. Subjects had to report whether they perceived a stimulus on each trial. Based on the detection rates of the pre-experiment, individual threshold contrasts for each of the 5 groups of stimuli were estimated by the experimenter. Occasionally, some of the contrasts had to be readjusted after the first block of the main experiment because subjects reported that they almost never saw anything.

Stimuli were presented on a light gray background with a luminance of 51.6 cd/m^2^. The luminance of the stimuli was 48.5 cd/m^2^ on average (median = 49.5 cd/m^2^, SD = 1.25 cd/m^2^, range = 46.5 – 51 cd/m^2^). The size of the stimuli was approximately 2.5 degrees of visual angle. Prior to the stimulus a fixation cross was presented. The size of the fixation cross was 0.35 degrees of visual angle and its luminance was 11.4 cd/m^2^. The response screen contained the question “Did you see something?” in the Estonian language. The contrast of the text was also low (luminance of 24 cd/m^2^), in order not to disturb the adaption of the eyes for very low contrast stimuli.

### 2.3. Task and design

Subjects were seated in a dark room, 80 cm from the monitor (SUN CM751U; 1024×768 pixels; 100 Hz refresh rate). Each session began with a short pre-experiment to determine the appropriate threshold contrasts for each subject (see 2.2 for more information), followed by the main experiment. The main experiment comprised 770 trials in total. Each of the 70 stimuli was presented 10 times. There were also 70 catch trials where no stimulus was presented. The order of the trials was fully randomized. Each trial began with the presentation of a fixation cross in the middle of the screen for 500 ms. The fixation cross was followed by a blank screen for 75-1250 ms. Then the stimulus was presented in the middle of the screen for one frame, that is or 10 ms, followed again by the blank screen. After 1s the response screen appeared.

Subjects were instructed to fixate on the cross in the middle of the screen, not to blink until the response screen had appeared, and then to report via button press on a standard keyboard whether they perceived a stimulus on a given trial or not. Seen and unseen responses were given with different hands, but the designated hands were balanced across subjects. There was a break after every 154 trials.

### 2.4. EEG recording and preprocessing

A Nexstim eXimia EEG-system with 60 carbon electrodes cap (Nexstim Ltd, Helsinki, Finland) was used. All 60 electrodes of the extended 10-20 system were prepared for recording. The reference electrode was placed on the forehead, slightly to the right. The impedance at all electrodes was kept below 15 KΩ. The EEG signal were sampled at 1450 Hz and amplified with a gain of 2000. The bandwidth of the signal was ca. 0.1 – 350 Hz. The horizontal electrooculogram (HEOG) was recorded in addition to the EEG.

EEG data was preprocessed with Fieldtrip (http://fieldtrip.fcdonders.nl; version 01-01-2013). Data were epoched around stimulus onset (-500 to +700 ms) and re-referenced to the average reference. Epochs were baseline corrected with a 100 ms time period before stimulus onset. All epochs were inspected manually for artifacts. Epochs containing blinks, eye movements, strong muscle activity or other artifacts were removed from the data. Noisy signals were interpolated with the nearest neighbor method. On average 2.6% of the data was interpolated (median = 2.6%, SD = 1.8%, range = 0.1 – 6.3 %). Data were filtered with a 30 Hz low-pass zero phase shift Butterworth filter.

### 2.5. Data analysis

The behavioral analysis was carried out with the R programming language (http://www.r-project.org/; version 3.1.0). EEG data was analyzed with Fieldtrip as well as with R.

#### 2.5.1 Behavioral analysis

As contrasts had to be readjusted occasionally during the main experiment (see 2.2), detection rate also varied in accordance with the different levels of contrast. In order to eliminate this accountable variance from the behavioral results only those contrast levels are considered which comprise the most trials. Thus, 93.3% of all available trials are considered (SD over subjects = 9%; SD over types of stimuli = 2.8%). Results are comparable, however, when all available trials are considered.

#### 2.5.2 Trial matching procedure

In order to find reliable markers of conscious perception and to characterize their timing (or variability thereof) 100 different matched sets of seen and unseen trials were constructed per subject. Each of these sets was composed as follows. For every stimulus on every contrast level (in case the contrast for this stimulus had to be readjusted after the first block of the main experiment) an equal number of seen and unseen trials were included in the respective conditions. In case there were only seen or only unseen trials in a given combination of stimulus and contrast level no trials were selected for either of the conditions. In case the number of seen and unseen trials in a given combination of stimulus and contrast level was unequal, all trials from the less numerous condition and an equal amount of randomly selected trials from the other condition were selected. This random selection of subsets was repeated 100 times for each subject. As a result both the seen and the unseen condition always comprised an equal number of trials for each subject on each iteration of the set matching procedure (m = 122, median = 123, SD = 31, range = 62 to 177). Note that stimulus content also remained identical for both conditions on every iteration.

After the trial matching procedure the seen and unseen conditions comprised 9.6 different types of stimuli (median = 10, SD = 0.6, range = 8 – 10) and 51.1 different individual stimuli (median = 52.5, SD = 8.2, range = 31 – 66) on 1.04 different contrast levels (median = 1, SD = 0.1, range = 1 – 1.4), on average. For all comparisons between the unseen and the catch condition all correctly rejected catch trials and the same sets of unseen trials were used. The catch condition comprised 59 trials on average (median = 60, SD = 5.6, range = 49 – 68).

#### 2.5.3 Cluster permutation tests

Differences between conditions were analyzed with nonparametric cluster permutation tests as implemented in Fieldtrip (Maris & Oostenveld, 2007). After averaging the single trials per condition data points (electrode-time pairs) were compared via dependent samples t-tests. Empirical distributions were created using 10 000 random permutations of the data. The maximal sum of t-values belonging to each cluster was used as the test statistic. Both the entry level for single samples into clusters and the significance threshold for clusters were set at .025. Only clusters lasting longer than 15 ms were considered significant. If not specified otherwise, cluster onsets and offsets were defined as the first/last time points when more than 4 neighboring channels showed significant differences between conditions.

#### 2.5.4 Denoising single trials

Based on results from a first set of cluster permutation tests representative channels of significant clusters were selected for denoising the single trial traces. The channels were Fcz, C1, Cz, C2, C4, CP1, Cpz, CP2, Pz for the P300 component and TP9, TP7, TP10, TP8, P10, P9, O1, Oz, O2, Iz for the N200 component. These channels were the first to show significant differences between conditions.

All available seen, unseen and catch trials were denoised together via an algorithm using wavelet decomposition (Quian Quiroga, 2000). This method allows the reconstruction of ERP components on the single trial level. The signal is first decomposed into different wavelets and subsequently reconstructed using only those wavelet coefficients that are relevant for the component of interest. Two different sets of wavelet coefficients were used for the reconstruction of the P300 and the N200, because the electrodes belonging to each cluster did not overlap. Nonetheless, the same sets of coefficients were used for each subject and each electrode within a given cluster.

#### 2.5.5 Correlation tests

After denoising the single trials (see 2.5.4) cluster permutation tests on the same sets of matched trials (see 2.5.3 and 2.5.2) were repeated to investigate significant differences between the seen and the unseen condition for the denoised data. To explain the variance in cluster onset latencies single trial parameters of the relevant components (N200 and P300) were correlated with the respective cluster onset latencies over all 100 iterations.

First, single trial parameters were extracted from the time period of observed variance in the onset latencies of significant clusters. For each trial, the positive peak between 151 - 268 ms was identified on each of the 9 reliable denoised electrodes belonging to the P300 cluster. Similarly, negative peaks were identified between 191 – 232 ms for the 2 reliable denoised electrodes (TP7 and P9) belonging to the N200 cluster. Both peak amplitude and peak latency were noted. These values were averaged for each subject on each of the 100 iterations. In addition to mean peak amplitude and mean peak latency, the standard deviation of peak latency was also computed for each subject on each iteration. These parameters were computed separately for the seen and the unseen condition. Finally, the 6 parameters (mean peak amplitude, mean peak latency and standard deviation of peak latency for both the seen and the unseen condition) were averaged over electrodes and subjects. Thus, a grand average of all 6 parameters for the N200 and the P300 per iteration was obtained. The grand averages were then correlated with the respective onset latencies of the N200 and the P300 clusters as obtained from the 100 permutation tests.

In addition to the 12 correlation tests described above 4 extra correlation test were carried out between averaged ERP parameters and cluster onset latencies. For these tests denoised single trial data was first averaged for each electrode per condition. Then, peak amplitude and peak latency of N200/P300 (depending on the electrode) was noted for the seen and the unseen condition. Finally, these values were averaged over electrodes and over subjects and correlation tests were carried out with the onset latencies of the respective clusters. All the p-values (n = 16) were corrected for multiple comparisons with the Holm-Bonferroni method.

## 3. Results

### 3.1 Behavioral results

The false alarm rate in our study was quite low considering the very faint stimulation. The mean percentage of seen reports for catch trials was 4.2% (median = 2.9%, SD = 4.5%, range = 0 – 17%). Mean detection rate over all stimulus types was 51% (median = 48.6%, SD = 13.8%). The high variance in detection rate stems from the fact that contrasts were estimated separately for different types of stimuli. For several subjects, threshold contrast could not be identified equally well for all stimulus types and detection rates were therefore not always clustered evenly around the mean. Table 1 contains detection rates for all stimulus types separately. Figure 2 depicts detection rates for all exemplars within the different types of stimuli.

**Table 1.** Means and standard deviations for the detection rates over subjects per stimulus type.

| Type | M | SD |
| --- | --- | --- |
| 1. animated solid forms | 0.48 | 0.36 |
| 2. animated line-drawing | 0.49 | 0.20 |
| 3. natural solid forms | 0.47 | 0.38 |
| 4. natural line-drawing | 0.36 | 0.20 |
| 5. faces (line-drawings) | 0.49 | 0.20 |
| 6. graphical solid forms | 0.39 | 0.29 |
| 7. graphical line-drawing | 0.57 | 0.26 |
| 8. text (solid forms) | 0.87 | 0.19 |
| 9. man-made solid forms | 0.47 | 0.37 |
| 10. man-made line-drawing | 0.50 | 0.20 |

**Figure 2.**
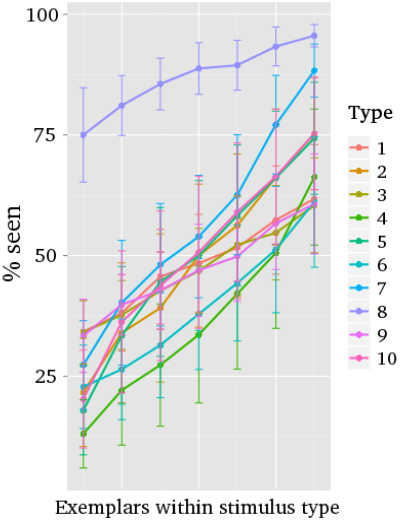
Variability in detection rates for exemplars within each stimulus type. Vertical lines represent standard errors. The stimulus type numbers correspond to the numbers in table 1.

As can be seen from Table 1, detection rates are considerably higher for text stimuli compared to other types of stimuli. This was due to the fact that for 12 out of 18 subjects no threshold contrast could be identified for text stimuli. Depending on the contrast, subjects either perceived close to none of the text stimuli or almost all of them. For those subjects the higher contrast level was selected and this pushed the mean detection rate up. For the other nine stimulus types threshold contrasts could be identified more successfully, but there was still variance between individual exemplars. Note, however, that the differences in detection rates were not systematic across exemplars. Some exemplars were perceived above average by some subjects and below average by others. Thus, in figure 2 exemplars are ordered according to their detection rate for each subject separately and it can be observed that the numbers do not necessarily refer to the same exemplars across all subjects.

### 3.2 EEG markers of conscious perception

It is evident form the behavioral results that the percentage of successfully perceived stimuli varies considerably between different stimulus types and even between single exemplars within a stimulus type. This, however, is not a problem for our present study. We are interested in the general markers of conscious perception (gmNCC in short). Such markers should not be affected by stimulus content variability. On the contrary, variance between stimuli can only strengthen any conclusions drawn from the results.

To study the onset latency of the gmNCC and its variability 100 different matched sets of trials with and without conscious perception were constructed (see *2.5.2 Trial matching procedure*). Consequently, a separate cluster permutation test was performed for each of the 100 sets (see *2.5.3 Cluster permutation tests*). If the results of these cluster permutation tests should turn out identical on every iteration then one could conclude that the gmNCC are highly reliable for the present study and that their timing remains constant.

However, the results demonstrate that there is considerable variability in the occurrence and timing of significant differences between the seen and unseen condition. Importantly, this variability is not caused by physical differences of the 100 iterations, as stimulus content remained identical for both conditions for each of the 100 iterations. Figure 3 shows the results of a representative permutation test. The most reliable difference between the seen and unseen condition was a positive cluster on central electrodes. This cluster was significant on every iteration, but the onset latency of statistical significance varied. Figure 4 contains a histogram of all observed onsets of the positive central cluster. It is obvious that there are two prominent periods of onset. Mean latency of the first onset period was 143 ms after stimulus presentation (median = 143, SD = 6 ms, range = 128 - 157). Mean latency of the second onset period was 193 ms (median = 190, SD = 13 ms, range = 166 - 223). This cluster was always significant until the end of the tested time period, i.e. 500 ms.

**Figure 3.**
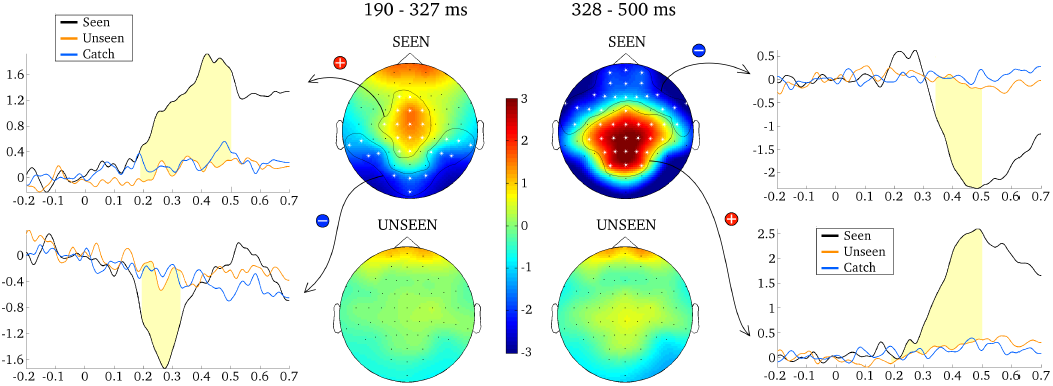
Results from one representative cluster permutation test. Topographies for the seen and the unseen condition are averaged over 190 – 327 ms (left) and 328 – 500 ms (right). ERP’s are shown for significant clusters, averaged over all electrodes belonging to each respective cluster (as indicated by white asterisks). Time periods where the seen and the unseen condition are significantly different from each other are colored light yellow.

**Figure 4.**
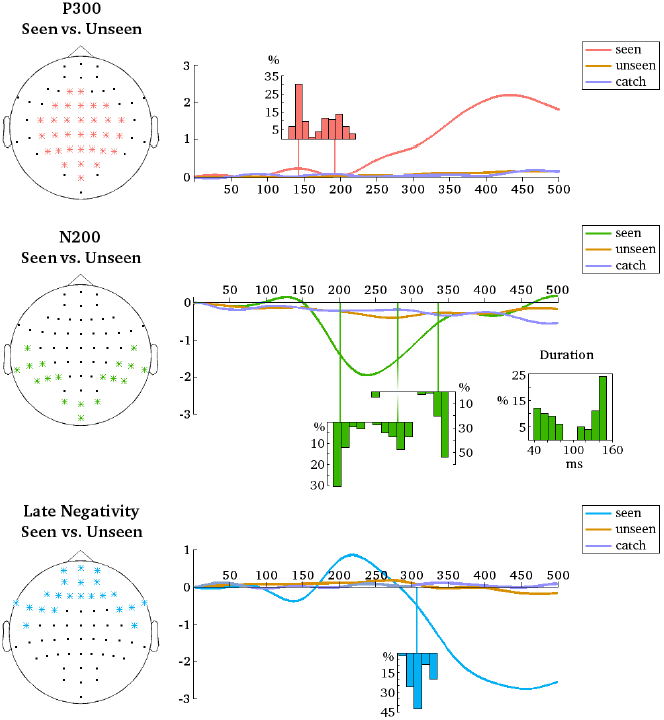
ERP’s are averaged over the indicated electrodes (left). These are all the electrodes that belonged to the respective clusters (P300, N200 or late negativity) for at least 1 of the 100 different permutation tests. Histograms depict the distributions of cluster onset times over the 100 permutation tests. For N200 there is also a distribution of cluster offset times and of cluster duration. Note that the distributions align with the time axes (in ms).

The positive central cluster constitutes the most reliable difference between the seen and the unseen condition because it was significant on every iteration. Furthermore, it was the earliest significant cluster on 91% of all iterations. There were, however, also two significant negative clusters. The more reliable of the two was a fronto-temporal cluster (Figure 3). Like the positive central cluster, this negative cluster was significant on all 100 iterations. The mean onset latency of statistical significance for this negative cluster was 307 ms (median = 309, SD = 11 ms, range = 283 - 334 ms, see also figure 4). Again, this cluster was always significant until the end of the tested time period.

The least reliable cluster was a negative occipito-temporal cluster, which was significant on 81% of the iterations (Figure 3). The onset of this negative cluster preceded the onset of the positive central cluster on 9% of the iterations. As for the positive central cluster, there are two prominent periods of onset for the negative occipito-temporal cluster. Mean latency of the first onset period was 203 ms (median = 199 ms; SD = 9 ms, range = 192 - 230 ms). Mean latency of the second onset period was 281 ms (median = 281 ms; SD = 12 ms, range = 257 - 301 ms). The duration of the negative occipito-temporal cluster was also divided into two groups. The first group lasted 59 ms on average (median = 57 ms, SD = 12 ms, range = 40 - 82 ms). The second group lasted 137 ms on average (median = 142 ms, SD = 12 ms, range = 108 - 150 ms). The mean offset of statistical significance was at 336 ms (median = 341 ms, SD = 20 ms, range = 245 - 350 ms). Figure 4 contains histograms of the distributions over all iterations.

To test if the above described components are uniquely associated with the seen condition we proceeded by comparing the unseen condition to the catch condition. A corresponding cluster permutation test did not yield any significant differences. Thus, it seems that a condition where the subject did not perceive a stimulus and a condition where there really was no stimulus are indistinguishable in our present dataset.

The first goal of the current study was to find general markers of conscious perception (gmNCC). The hitherto results suggest two such markers for our dataset. The most prominent marker is a positive cluster of central electrodes. We will call this component P300. A less reliable but nonetheless noteworthy marker is a negative cluster on occipito-temporal electrodes. We will call it N200. The onset and topography of the third late negative cluster on fronto-temporal electrodes suggest that it is a consequence of conscious perception (Aru et al., 2012). It is of course also possible that this late negativity is a secondary stage of the conscious experience of a stimulus reflecting its holding or maintenance in working memory. We leave this problem out of the scope of the present article, however, and will not concentrate on this component any further.

The second goal of the present study was to examine the timing of gmNCC as closely as possible. The above described results would suggest that the timing of gmNCC is highly variable, ranging over 100 ms depending on which trials are included in the comparisons. It is important to keep in mind, however, that the presently described variability in onset latencies could have occurred due to several reasons and the results do not allow any strong conclusions with regard to the actual gmNCC onset latencies (see 1. Introduction). Thus, in order to zero in on a better estimate of latency for the potential gmNCC one must first try to identify the sources of the observed variability in onsets of statistical significance. One would have to investigate how much the true latency differences of gmNCC contribute to the estimate of the gmNCC onset latency as compared to other possible sources such as noise.

### 3.3 gmNCC onset variability explained by noise

We have so far observed differing onsets of statistically significant effects for the N200 and the P300 components depending on which trials are included in the comparisons. It follows that some variables characterizing single trials are responsible for the varying results. We will now try to identify these variables. As stated above, both the signal and the noise profiles of the single trials are potentially involved. Thus, the first objective was to eliminate the contribution of noise as much as possible. We wanted to know how the results presented above are changed when the possible effect of noise is diminished. Keep in mind that this is also done in order to arrive at the best possible estimate of latency for the gmNCC.

Nine representative channels of P300 and ten representative channels of the N200 were selected. The single trial traces on these channels were denoised via wavelets (see 2.5.4). Then the 100 cluster permutation tests were repeated on the same sets of seen and unseen trials as for the undenoised data. After denoising, the onset latency of statistically significant differences again showed considerable variance, albeit with some important differences. The previously observed early period of P300 onsets was effectively not present. Only two from the 100 iterations resulted in P300 onsets earlier than 160 ms. The mean onset latency for the new results was 232 ms (median = 231 ms, SD = 17 ms, range = 151 - 268 ms). Figure 5 contains the distribution of all onset latencies after denoising the data. Again, the P300 always remained significant until the end of the tested time period.

**Figure 5.**
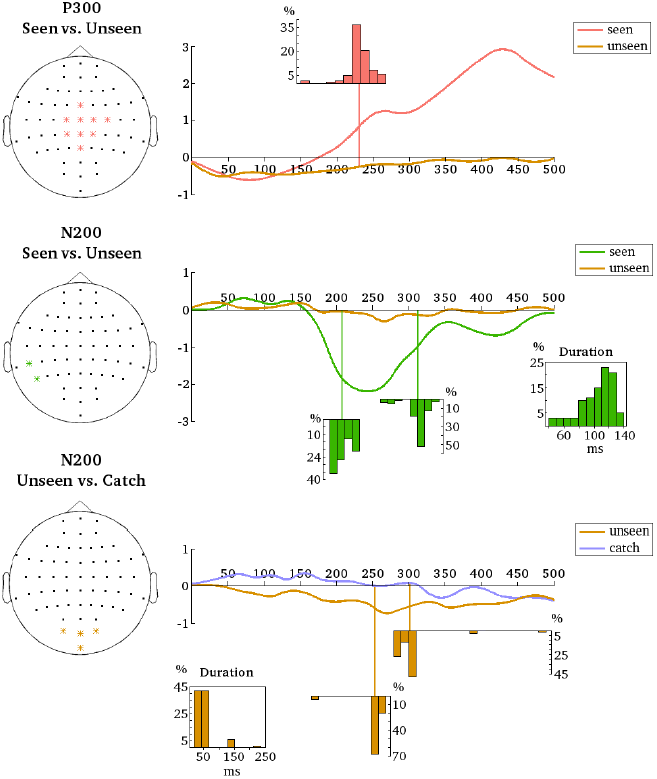
Denoised data is averaged over the indicated electrodes (left). These are all the electrodes that are most representative (i.e. most reliable) for the respective clusters (P300 and N200 for the seen-unseen comparisons; N200 for the unseen-catch comparisons). Histograms depict the distributions of cluster onset times, cluster offset times and cluster durations over the 100 different permutation tests. The distributions align with the time axes (in ms).

Results also changed for the N200. After denoising it was observed that N200 was smaller in topographical size. Only 2 temporo-parietal channels showed reliable effects. We nonetheless decided to go on with the analyses considering clusters starting from 2 channels as significant. For the new results N200 was significant on 97% of the iterations and included on average 3.3 channels (median = 2, SD = 2, max = 9). The mean onset latency was 207.9 ms (median = 203.4 ms, SD = 12.3 ms, range = 191 − 232.4 ms). The mean offset latency was 312.7 ms (median = 316.6 ms, SD = 14.8 ms, range = 261.4 − 342.1 ms). Thus, the mean duration of the N200 was 104.9 ms (median = 111 ms, SD = 22.6 ms, range = 39.3 − 141.4 ms). Figure 5 contains histograms of the respective distributions. Note that the N200 onset and duration displayed a highly negative correlation (r = −0.8, t(95) = −12.95, p = 2.2e-16). The onset of the N200 and the onset of the P300 were positively correlated (r = 0.23, t(95) = 2.27, p = 0.025). The duration of the N200 and the onset of the P300 were slightly negatively correlated (r = −0.19, t(95) = −1.9, p = 0.06).

In addition to the comparisons between the seen and the unseen condition a separate group of a hundred permutation tests comparing the unseen condition to the catch condition were performed. Denoised data were analysed from the same groups of channels that were used for the seen-unseen comparisons. The goal was to examine if the N200 and the P300 are uniquely associated with the seen condition without too much contribution from prerequisites of NCC (Aru et al., 2012). Recall that no corresponding differences for the undenoised data were found, but perhaps the removal of noise will bring to light some subliminal processing of the stimulus in the unseen condition that was previously missed.

As for the undenoised data, there were no significant differences between the unseen and catch conditions on the central P300 channels. Thus, the P300 seems indeed to be only associated with the seen condition. In a situation where the stimulus was not perceived no P300 can be identified. The same is not quite true for N200, however. The hundred cluster permutation tests comparing the unseen condition to the catch condition on occipito-temporal channels revealed a small but quite consistent occipitally confined negative cluster. Note that these are not the same channels that were most reliable in the seen-unseen comparison. The negative occipital cluster was significant on 91% of the iterations and included 3 electrodes on average (median = 3, SD = 0.21, max = 4). The mean onset latency of statistical significance was 253.7 ms (median = 255.9 ms, SD = 16.5 ms, range = 165.5 − 268 ms). The mean offset latency was 301.6 ms (median = 300 ms, SD = 27.7 ms, range = 278.6 − 491 ms). Thus, the mean duration of the occipital negative cluster was 47.9 ms (median = 42.8 ms, SD = 33 ms, range = 20 − 235.2 ms). Figure 5 contains histograms of the respective distributions.

It is thus evident that a negative cluster on occipital electrodes can reliably differentiate the unseen condition from the catch condition around 250 ms after stimulus onset. Still, based on consistent differences in topography and latency, one can be fairly confident that it is not the same component as the N200 from the seen-unseen comparison. We therefore conclude that the N200 on a small cluster of left temporo-parietal channels is also uniquely associated with the seen condition. This does not necessarily mean that the same neural mechanisms may not be involved in pre-conscious and conscious processing. They can be the same, but the latency of becoming involved and the level of expression of activity are different. Foremost, ERPs are signatures of neural activity rather than neural structure.

After having removed noise from the data the results did indeed get more homogeneous. It seems that random summation of noise was the reason behind some of the variance observed in the results from section 3.2. But even if the data was effectively denoised some variability in the onset of statistically significant differences between the seen and the unseen condition still remained. One can therefore already conclude that it was not only the noise profile of the single trials that was responsible for the variability of the onset latency of the gmNCC presented in section 3.2. One might suspect that some parameters of the signal profile (e.g. latency/amplitude of the components) are also involved in the observed variance. Thus, we next try to identify the parameters of the denoised single trials that determine the onset of significant differences between the seen and the unseen condition. Through that analysis we also hope to arrive at the best possible estimate of latency for the gmNCC.

### 3.4 gmNCC onset variability explained by single trial parameters

Having the list of 100 varying onset times of statistically significant differences between the seen and the unseen condition one can ask what is different between those denoised trials that constitute the respective sets for each iteration. To answer this question some key parameters of the single trial N200/P300 were extracted from the respective time periods of observed variability in cluster onset latency. Then the hundred different cluster onset latencies for N200/P300 were correlated with grand averages of the extracted parameters from those denoised trials that were included in the respective iterations. These parameters were mean peak amplitude, mean peak latency and mean latency variance for the seen and the unseen trials (see 2.5.5 for more details).

It is important to note that we are presently not analyzing the peaks of the N200 and the P300 components. Because we are interested in the time period of cluster onsets we cannot hope to accurately capture the peaks of the corresponding components in that time window. Our aim is somewhat different. We are trying to understand what happens in the single trials at the time when variance is observed between the 100 iterations. We are trying to do this by looking at maximal activity in that time window. Because we already have our significant cluster permutation test results, we know that some variables must exist that are responsible for the differences in the results. We are now simply taking our analysis one step further by trying to identify these variables. Obviously, the results of our cluster permutation tests are based on reliable differences in amplitude, but as we have already outlined above (see also Navajas et al., 2013) differences in amplitude can also occur due to monotonic shifts in latency or due to differences in latency variance. That is the reason why we have included these three parameters (amplitude, latency and latency variance) in our current correlation tests.

Table 2 contains the results of all conducted correlation tests and figure 6 illustrates them. The cluster onset times of P300 correlated significantly neither with mean peak latency of the seen trials nor with mean peak latency of the unseen trials. The respective correlations with mean latency variance were also not significant. There was a moderately significant correlation with mean peak amplitude for seen trials, but the most significant correlation was found with mean peak amplitude for unseen trials. Results were very similar for N200. The cluster onset times of N200 did not correlate significantly with mean peak latency or mean latency variance for the seen or for the unseen trials. The correlations with mean peak amplitudes of the seen and the unseen trials were again significant. Thus it seems that the varying onset times of the two general markers of conscious perception are first and foremost explained by amplitude variability in the unseen trials, but amplitude variability in the seen trials has an effect as well.

**Table 2.** Correlations between mean amplitude, mean latency or mean latency standard deviation of single trial N200 and P300 responses with cluster onset times (see materials and methods for the construction of different sets of trials and the corresponding cluster permutation tests). Correlation test are carried out separately for the seen and the unseen trials. All p-values are Holm-Bonferroni corrected for 16 tests in total (see also table SI 3.1).

|  |  | **Mean Amplitude** | **Mean Latency** | **Mean Latency SD** |
| --- | --- | --- | --- | --- |
| **N200** | SEEN | $r = 0.34, t = 3.49, p = 0.009$ | $r = -0.02, t = -0.18, p = 1.0$ | $r = 0.22, t = 2.25, p = 0.21$ |
| | UNSEEN | $r = -0.48, t = -5.4, p = 7e-06$ | $r = 0.01, t = 0.09, p = 1.0$ | $r = -0.23, t = -2.26, p = 0.21$ |
| **P300** | SEEN | $r = -0.3, t = -3.07, p = 0.031$ | $r = -0.05, t = -0.52, p = 1.0$ | $r = 0.04, t = 0.43, p = 1.0$ |
| | UNSEEN | $r = 0.5, t = 5.67, p = 2.2e-06$ | $r = 0.15, t = 1.54, p = 0.76$ | $r = -0.02, t = -0.21, p = 1.0$ |

**Figure 6.**
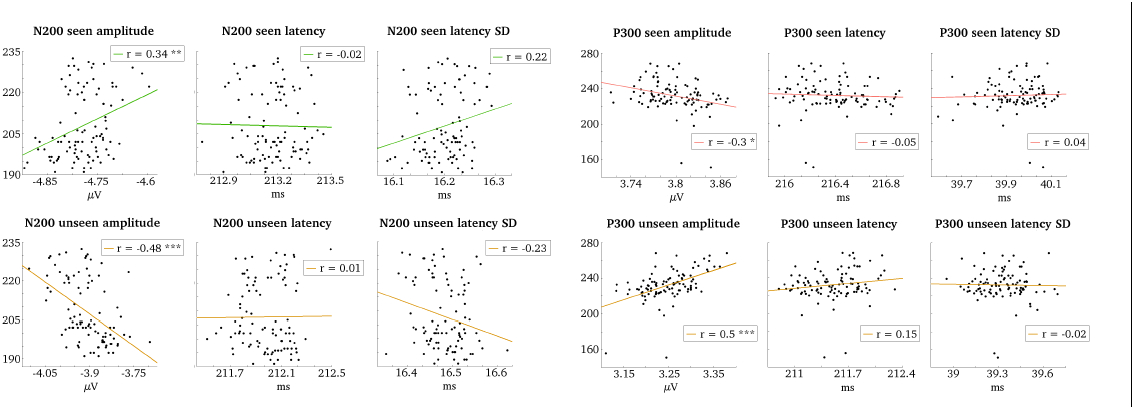
Correlations between grand averages of single trial N200/P300 parameters (amplitude, latency and standard deviation of latency) and N200/P300 onset times (indicated in ms on the y-axes). Correlation tests are carried out separately for seen and unseen trials. P-values < 0.05 are indicated with *. P-values < 0.01 are indicated with **. P-values < 0.001 are indicated with ***.

To exclude any possible confounds with latency variance and to demonstrate more convincingly the relevance of the amplitude parameter for the observed variability in cluster onset times, the above analysis was repeated by first averaging single trials and then extracting peak amplitude. The correlation between mean peak amplitude of the averaged seen trials and the P300 cluster onset times was not significant, but the correlation for mean peak amplitude of the averaged unseen trials was again highly significant. Similarly, the correlation of mean peak amplitude with N200 cluster onset times for the seen trials was only marginally significant. The same correlation for unseen trials was again highly significant. Table 3 contains the results for these correlation tests with averaged data and figure 7 illustrates them.

**Table 3.** Correlations between mean N200 or P300 amplitudes (after averaging the single trial responses per subject) with cluster onset times (see materials and methods for more information). Correlation test are carried out separately for the seen and the unseen trials. All p-values are Holm-Bonferroni corrected for 16 tests in total (see also table 2).

| Mean amplitude of averaged trials |  |  |  |  |  |
| --- | --- | --- | --- | --- | --- |
| **N200** | SEEN | $r = 0.28, t = 2.9, p = 0.049$ | **P300** | SEEN | $r = -0.26, t = -2.63, p = 0.089$ |
| | UNSEEN | $r = -0.47, t = -5.14, p = 1.9e-05$ | | UNSEEN | $r = 0.53, t = 6.17, p = 2.5e-07$ |

**Figure 7.**
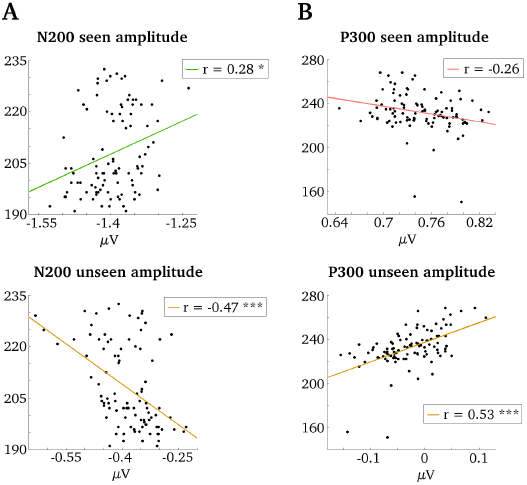
Correlations between grand averages of N200 and P300 amplitudes (after averaging the single trial responses per subject) and the respective cluster onset times (indicated in ms on the y-axes). Correlation tests are carried out separately for seen and unseen trials. P-values < 0.05 are indicated with *. P-values < 0.001 are indicated with ***.

We conclude that it is predominantly the mean amplitude of unseen trials that defines the onset of significant differences between the seen and the unseen condition and therefore the onset of the gmNCC. Higher mean amplitude on central channels during the 151 – 268 ms period for unseen trials is associated with later onsets for P300 in the seen-unseen comparisons. Similarly, more negative mean amplitude on left temporo-parietal channels during the 191 – 232 ms period for unseen trials is associated with later onsets for N200 in the seen-unseen comparisons. If the range of mean amplitude values for the seen and the unseen trials in figure 6 are compared, it can be noticed that mean amplitude of the unseen trials varies over a wider range than mean amplitude of the seen trials. Thus, it is not surprising that this variability is reflected in the onset times of significant differences between the seen and the unseen condition. However, cluster onset times are also influenced by variability in mean amplitude of the seen trials. Lower mean amplitude on central channels during the 151 – 268 ms period for seen trials is associated with later onsets for P300 and less negative mean amplitude on left temporo-parietal channels during the 191 – 232 ms period for seen trials is associated with later onsets for N200.

Most importantly, there is no evident connection between cluster onset times and latency parameters. Neither mean latency nor mean latency variance of N200 and P300 were reliably correlated with the onset times of the respective significant clusters. And indeed, if one takes a look at the distributions of mean P300 and mean N200 latencies in figure 6, respectively, one can observe that the variability is very small in absolute numbers. It seems that the mean latencies of the two gmNCC are very similar across the different sets of trials. The distributions of mean latency variance for P300 and N200 in figure 6 make it clear that peak latency shifts considerably over single trials, but mean latency variance is again very similar across the 100 different sets of trials. We are therefore able to conclude that the mean latency of the gmNCC does not shift in time as much as the results from seen-unseen comparisons would suggest. Rather, it seems that mean latency of gmNCC stays fairly consistent, but the gmNCC are expressed in different strengths (i.e. have different amplitudes) for the different sets of trials.

## 4. General discussion

The first goal of the present experiment was to find general markers of NCC (gmNCC), that is - markers of conscious perception common to all cases where the stimuli were perceived and absent for all cases where the stimuli were not perceived. The second goal was to study how much the timing of these gmNCC varies within one experiment. A heterogeneous visual stimulus set was presented at a near-threshold contrast. Thus, the applied paradigm was designed to reduce the influence of stimulus predictability and categorical and individual specificity. 100 different subsets of the resulting seen and unseen trials were contrasted to identify the gmNCC and to study the variability in their timing. Results indicate two gmNCC for our paradigm. First, a more prominent positive cluster on central electrodes (P300) and second a negative cluster on left temporo-parietal electrodes (N200). Furthermore, we observed that there is considerable variability of the onsets of N200 and P300 even within one single study.

The P300 component is a well known marker of conscious perception. It has been found in almost all electrophysiological studies investigating the ERP-correlates of consciousness. Only when the same experimental stimuli are presented repeatedly (Koivisto & Revonsuo, 2008; Sekar et al. 2013) or when one has prior knowledge about the presented stimulus (Melloni et al., 2011) does the P300 not occur as a difference between trials with and without conscious perception. It can be argued that in both cases the upcoming stimulus is fully determined by the prior event. As P300 might reflect updating of working memory (WM) (Polich, 2007), which is arguably not needed when the very same stimuli are already encoded in WM, P300 is not a marker of conscious perception under such experimental conditions (Melloni et al., 2011). For the present study stimuli were deliberately unpredictable. In the light of the argumentation presented above it is not surprising that the P300 is a prominent gmNCC in our dataset.

The N200 has also been found as a marker of conscious perception, but not as often as the P300. In many studies the N200 is not reliably different between conditions with and without conscious awareness (e.g. Sergent et al., 2005; Del Cul et al., 2007). The present results offer an explanation for these varying results. As the reliability of this marker of conscious perception depends on which single trials are included in the seen as well as the unseen condition it is possible that previous studies have simply missed it. This possibility has also been noted by Del Cul et al. (2007). Nonetheless, the present results are different as far as there really is no clear N200 component present in the unseen condition of our study. Studies using stronger stimuli find a well pronounced N200 which is not different between conditions (e.g. Sergent et al., 2005; Del Cul et al., 2007). Thus, one might argue that the N200 reflects a process preceding the NCC proper.

Yet, it seems for the undenoised data that the average onset of P300 is somewhat earlier than the average onset of N200. This would be in conflict with the view that N200 reflects a pre-conscious process prior to the NCC proper, which is P300. Another interesting observation is that both components show two periods of onset for the undenoised data. One explanation for these results is that the abnormally distributed results are due to a confounding signal in the measurements and are actually not a property of the gmNCC per se. The current results favor this explanation because after denoising the relevant single trial data, the divided periods of onset disappear. After denoising both components are still reliably associated with conscious perception, but they show one fairly similar period of onset which falls around 200 ms after stimulus presentation.

Despite the more coherent results for denoised data the variance in gmNCC onset latencies still remains. Different onset latencies of NCCs observed in different studies can be parsimoniously explained with differences in stimulus material and tasks (Dehaene & Changeux, 2011). However, here we observed large variability of NCC onsets even with the very same stimulus material and task within one study. The variability was evident when simply considering different trials for the compared conditions. This implies that when random subsets of trials are selected, the onset and duration of the NCC depends on the particular random subset of trials and the onset of the NCC can vary as much as 100 ms dependent on this random selection.

Interestingly, we were able to show that the variance of the onset latency of the markers of conscious perception can be first and foremost attributed to amplitude variance in the unseen condition. This is not directly relevant for the gmNCC because, compared to the seen condition, there are no equivalent components in the unseen condition as shown by comparisons with the catch condition. However, this result shows that the trials from the unseen condition have an important influence on the timing of gmNCC. In other words, the specific trials of the unseen condition influence the first time point and the time intervals where significant differences between trials with and without conscious perception are observed. Amplitude variance in the seen condition is also associated with the varying gmNCC onsets, albeit to a lesser extent. In contrast to amplitude, N200/P300 latency and latency variance does not seem to have an influence on gmNCC onset latency. Overall, the mean onset latency of the two markers is fairly consistent independent of their amplitude.

Our goal was to arrive at the best possible latency estimate for our two gmNCCs. The presently reported results bring us closer to an informed answer. We now know that it is not justified to only use the onset times of significant differences between the seen and the unseen condition for an estimate of gmNCC latencies. In fact, one can disregard a lot of the variance in cluster onset times because it is caused by irrelevant amplitude fluctuations in the unseen condition. Recall that N200 and P300 are uniquely associated with the seen condition. If similar components would be present in the unseen condition (which would make them relevant for the seen-unseen comparisons) one would have found more evidence for them in the unseen-catch comparisons. Still, even if it is known that the variability in cluster onset times is not indicative of variability in the latency of the gmNCC, it is not known which cluster onset value to take as the estimate of the gmNCC latency. Is the mean cluster onset time better than the earliest cluster onset time? Or is median cluster onset time the right choice? There really is no way of telling because one cannot fully disentangle the contribution of mean amplitude for the seen trials from the contribution of mean amplitude for the unseen trials. Both vary and both have an effect on the onset time of significant differences.

With this in mind, the best and most reliable estimate of the latencies of N200 and P300 should not be based solely on results from the cluster permutation tests, but also on original latency values for the seen trials in the time window of significance onsets, because these seem to stay surprisingly homogeneous over different sets of trials. Presently, we will use mean peak latency to make the best estimates. For N200 latency the respective estimate is 213 ms (SD over subjects = 2.2 ms, range = 210 – 218 ms). For P300 latency the estimate is 216 ms (SD over subjects = 5.5 ms, range = 209 – 227 ms). Whether the components will be significant at these times in a given seen-unseen comparison depends a lot on mean amplitudes of the specific selection of the seen and unseen trials included in the comparison. But the mean peak latency of these gmNCC gives an idea of what is going on in the seen trials alone.

It is noteworthy that not only are the N200 and P300 components missing in the unseen condition, but there are really no clear ERP’s associated with the unseen condition at all. Ojanen et al. (2003) conducted an experiment with a similar paradigm to ours and they also did not find any clear ERP’s for the non-perceived stimuli. Thus, it seems that unconscious perceptual processes evoked by the stimuli are too faint or too unsystematic to be picked up via EEG. But when a closer look is taken at the available data (i.e. denoising the single trials) then it becomes evident that the unseen and the catch conditions can nevertheless be reliably different on occipital electrodes around 250 ms after stimulus presentation. This fact points at the possibility that the processes underlying conscious perception have a varying level or degree of expression which at a certain definite value specifies the threshold of conscious perception.

Although the same occipital electrodes can sometimes also show significant differences between the seen and the unseen condition, these are not the most reliable electrodes for the N200 of conscious visual perception. N200 was most reliable on left temporo-parietal electrodes in the present study. Thus, one additional possibility why some previous works have not found the N200 as a marker of conscious perception could be because it is mixed up with other posteriorly recorded components that have similar latencies, but are not necessarily associated with conscious perception.

In addition to the flat ERP of the unseen condition we also did not observe early EEG components in the seen condition (e.g. N100) for the present paradigm. Again, it is likely that these signals are too faint and/or unreliable for the low contrast stimuli used in the present study. This interpretation is backed up by a study of Sekar et al. (2013) who also used very weak stimulation and found very small post-stimulus brain response at 100 ms that did not differ between conditions. Thus, the present results confirm that such early responses do not seem to be markers of conscious perception.

What are the theoretical implications of the two gmNCC found in the study at hand? The P300 is most consistent with the theory of a global workspace consisting of multiple areas including frontal, parietal, and temporal cortices (Dehaene et al., 1998; 2003). We cannot say anything certain about the sources of our P300, but since it is a very well studied component one can be fairly confident that a similar multi-focal network is underlying the P300 of the present study.

In addition to the P300 we also find a more transient component with a similar onset latency – the N200. This gmNCC could be consistent with the visual awareness negativity (Koivisto & Revonsuo, 2003; Wilenius-Emet et al., 2004) concept and the idea of posterior local recurrent activity (Lamme & Roelfsema, 2000). Our N200 component occurs somewhat later and is less consistent than the usual N200 reported previously. This may be due to the faint stimulation. A similar explanation is offered by Sekar et al. (2013). The facts showing that ERP correlates of correct perception have been found at a shorter latency range exemplified by N100-150 (Bachmann, 1994) can be explained as a result of the considerably higher contrast/intensity of the stimuli used, which leads to the speed-up of awareness-related processing and shorter latencies of the negative ERP components reflecting this.

Taken together, our findings show that if a set of heterogeneous stimuli is used, whose identify cannot be predicted by the subject, the two widely reported correlates of consciousness – the N200 and P300 – are reliably observed. These two ERP components are the two general markers of conscious perception found in our study. However, the onset latencies of these components still showed large variability. Importantly, part of this variability can be attributed to the particular set of trials selected for the condition without conscious perception. These results indicate that any conclusions about the NCC onset timing that are based on data from a single study are likely to be misleading.

## Acknowledgments

We thank Dr. Lucia Melloni for her helpful advice during the preparation of the manuscript.

**Research reported in this paper is supported by Estonian Science Foundation, project SF0180027s12 (TSHPH0027) and Estonian Science Agency (IUT40-30).**

